# Replicated repurposing of an ancestral transcriptional complex in land plants

**DOI:** 10.1101/2025.04.25.650595

**Authors:** Thea E. Kongsted, Facundo Romani, Chiara A. Airoldi, Jim Haseloff, Beverley J. Glover

## Abstract

Transcriptional complexes with a common composition regulate the production of flavonoid pigments, trichomes, root hairs and other epidermal traits in seed plants. These complexes are composed of transcription factors from the MYB and basic helix-loop-helix (bHLH) families along with a tryptophan-aspartate repeat (WDR) scaffold protein (MBW complexes). The MYB member has been found to be the most pathway-specific component of the complex and modifications to these *MYB* genes are overrepresented in studies investigating the genetic basis of changes in pigmentation phenotypes across flowering plants. Here we investigated the orthologues of the MBW complex in a divergent lineage to understand its origin and evolution. We found evidence that these transcriptional complexes also form in the liverwort *Marchantia polymorpha*, indicating, together with an analysis of published gene family phylogenies, that they are ancestral to land plants. The functions of each of the two orthologous *MYB* genes, Mp*MYB14* and Mp*MYB02*, both depend on the single orthologous *bHLH* gene, Mp*bHLH12*. We could not assess the functional role of the *WDR* genes in *M. polymorpha*, due to low mutant recovery suspected to be caused by pleiotropic effects on viability. We propose that the two transcriptional complexes with alternative *MYB* paralogues in *M. polymorpha* represent an ancestral function, regulation of the flavonoid pathway, and a derived function, maturation of liverwort-specific oil bodies. Our findings imply a replicated pattern by which new complexes have evolved in independent land plant lineages, through duplication of the evolutionarily labile *MYB* member and co-option of its interaction partners.

## Introduction

Across taxa, some loci seem to be overrepresented in studies investigating the genetic basis of phenotypic differences. For instance, a few recurrent loci have been found to be associated with the evolution of diverse wing colour patterns across divergent lineages of butterflies and moths (Orteu and Jiggins, 2020). Systematic identification of such “hotspots” of evolution may give broader insights into how the developmental processes that give rise to phenotypes structure evolutionary change (Stern and Orgogozo, 2009). In the flowering plants, red, purple and blue colours are most commonly caused by anthocyanin pigments derived from the flavonoid pathway. The evolution of pigmentation phenotypes in both natural and cultivated populations has been found to show higher association with changes to the activating MYB transcription factors than any other component of this pathway (Sobel and Streisfeld, 2013; Wheeler *et al*., 2022; Marin-Recinos and Pucker, 2024). One such *MYB* gene (*COLOURED ALEURONE1*) was the first transcription factor identified in a plant and the site of integration when transposable elements were first discovered by Barbara McClintock in variegated *Zea mays* lines (Paz-Ares *et al*., 1987). These MYB proteins regulate the transcription of enzymes in the flavonoid biosynthetic pathway as part of higher-order transcriptional complexes known as MYB-bHLH-WDR (MBW) complexes. In the model flowering plant *Arabidopsis thaliana*, MBW complexes have also been identified that regulate other epidermal phenotypes, including the *de novo* patterning of trichomes on aerial organs (Zhang *et al*., 2003), the position-dependent spacing of both root hairs (Bernhardt *et al*., 2003) and hypocotyl stomata (Berger *et al*., 1998), and the differentiation of the seed coat (Baudry *et al*., 2004). In each case, the MYB member of the complex is pathway-specific (Ramsay and Glover, 2005). Overlapping subsets of MYB proteins form complexes with each of four bHLH proteins, which thus show several functions and partial redundancy (Zhang *et al*., 2003). Finally, a single WDR scaffold protein, TTG1, is required for the functions of all of these complexes (Walker *et al*., 1999). To further investigate evolutionary lability in these pathways, we set out to explore the origin and functional diversification of the MBW complexes.

So far, MBW complexes have been studied extensively in the angiosperms (reviewed in Ramsay and Glover, 2005) and one report found that they also regulate the flavonoid pathway in a gymnosperm (Nemesio-Gorriz *et al*., 2017). As such, their origin can be traced back at least as far as the stem lineage of seed plants. However, the participating gene families are found across eukaryotes (Roussel *et al*., 1979; Murre, McCaw and Baltimore, 1989; Smith *et al*., 1999) and as such, it is not clear when the complexes first evolved. In the following paragraphs, we introduce in turn what can be gleaned from published phylogenetic analyses about the origins of MBW complex formation in the MYB, bHLH and WDR gene families.

The ∼20 MYB proteins that participate in MBW complexes in *Arabidopsis thaliana* share a common motif in the R3 MYB repeat (Zimmermann *et al*., 2004, see also Grotewold *et al*., 2000). In a phylogenetic analysis of R2R3MYB genes in land plants and their closest relatives, the Zygnematophyceaen charophyte algae, Jiang & Rao (2020) found that this motif is restricted to, and broadly conserved across, a land plant-specific clade, named VIII-E in their streptophyte-level nomenclature. The most parsimonious scenario would appear to be an origin of the motif at the base of this clade, and thus with a duplication in the stem lineage of land plants, with several shallower secondary losses. In the *A. thaliana* nomenclature (Stracke, Werber and Weisshaar, 2001) VIII-E corresponds to four subgroups that form MBW complexes (4, 5, 6 and 15 (Zimmermann *et al*., 2004)) and three that do not (7 (act alone, see Mehrtens *et al*., 2005; Stracke *et al*., 2007), and the closely related 19 and 20 (19 form complexes with bHLH proteins belonging to subclass IIIe, see Qi *et al*., 2015). This correlates with presence/absence of the motif across these subgroups (Jiang and Rao, 2020). The authors found that the motif was present not only in sequences from seed plants, where MBW complexes have been reported, but also in sequences from other vascular plants, and in two sequences from the moss *Physcomitrium patens*, which belongs to the other monophyletic group of land plants, the bryophytes. Another bryophyte, the liverwort *Marchantia polymorpha* encodes two VIII-E MYB genes (Jiang and Rao, 2020), both of which show partial conservation of the motif, lacking two of the six conserved residues identified in *A. thaliana*. One of these genes, Mp*MYB14,* is known to activate the production of flavonoids, including red auronidin pigments (Albert *et al*., 2018; Berland *et al*., 2019) in response to different environmental challenges including mineral nutrient insufficiency (Albert *et al*., 2018), high light (Albert *et al*., 2018) and oomycete infection (Carella *et al*., 2019). The other, Mp*MYB02*, is required for the maturation of oil bodies (Kubo *et al*., 2018; Wang *et al*., 2023). Oil bodies are membrane-bound organelles which are synapomorphic to the liverwort lineage (Kanazawa *et al*., 2020; Romani *et al*., 2022). They accumulate cytotoxic terpenoids and phenylpropanoids, including liverwort-specific bis-bibenzyl compounds (Tanaka *et al*., 2016), which serve as a defense against herbivory (Romani *et al*., 2020). The limited conservation of the interaction motif in the R3 MYB repeats of the *M. polymorpha* orthologues has previously been taken to imply that they act independently of an MBW complex (Davies *et al*., 2019).

The bHLH proteins that participate in MBW complexes in seed plants all belong to subclass IIIf, while a member of subclass III(d+e) has been shown not to interact with their MYB partners (Zimmermann *et al*., 2004). Recently, we found that these two subclasses both derive from early land plant duplications, and that they are sister to one another (Kongsted and Glover, 2023). This suggests an origin of MBW participation for bHLH proteins coincident with that inferred above for MYB proteins, in the stem lineage of land plants. *Physcomitrium patens* encodes four IIIf bHLH proteins, while *M. polymorpha* encodes a single member, Mp*bHLH12* (Kongsted and Glover, 2023), the function of which has not yet been clearly determined (although see work by Arai *et al*., 2019).

For the scaffold protein AtTTG1, highly sequence-conserved putative orthologues can be identified across eukaryotes (De Vetten *et al*., 1997). The At*TTG1* gene derives from a duplication at the base of seed plants (Airoldi *et al*., 2019), which also gave rise to the progenitor of the Brassicaceae-level paralogues At*LWD1* and *2*, which redundantly regulate circadian rhythms (Wu, Wang and Wu, 2008). Three homologues have been identified in *M. polymorpha*, two of which (either Mp*WDR1* or Mp*WDR2*, but not Mp*WDR3*) can individually complement both epidermal and circadian defective phenotypes when heterologously expressed in an *A. thaliana ttg1 lwd1 lwd2* mutant, while the *A. thaliana* paralogues can not complement one another’s functions (Airoldi *et al*., 2019). This suggests that the seed plant duplication was followed by subfunctionalisation and that an ancestral TTG1-like protein in the land plants was capable of scaffolding MBW complexes (Airoldi *et al*., 2019).

Based on this synthesis of previous work, we hypothesised that the diverse MBW complexes characterised in seed plants derive from an ancestral MBW complex that originated in the stem lineage of land plants, associated with a new paralogue in the *bHLH* and *MYB* families, and the recruitment of a pre-existing WDR protein. To test this hypothesis, we asked whether orthologues from *M. polymorpha*, which diverged from the lineage leading to seed plants at the first bifurcation of crown group land plants, form transcriptional complexes. We found evidence that the *M. polymorpha* orthologues form complexes to regulate pigment biosynthesis and oil body maturation, although the functional role of the WDR scaffolds could not be confirmed. We propose that one complex in *M. polymorpha* retains the ancestral function, shared with seed plants and conserved since stem land plants, while another arose within the liverwort lineage. Notably, we infer that the origin of this new complex with a derived function mirrors what occurred independently in the angiosperm lineage.

## Results

### The two *Marchantia polymorpha* VIII-E MYB proteins can each form complexes with the single IIIf bHLH protein

As noted, a motif in the R3 repeat is conserved across those of the *A. thaliana* R2R3MYB proteins (e.g. AtWER, GL1, PAP1 and TT2 shown in Fig 1A) and single-repeat R3MYB proteins (e.g. AtCPC shown in Fig 1A) that form MBW complexes. This motif is not conserved in members of the closest paralogous clade, VIII-D (e.g. AtMIXTA and MpSBG9 shown). The two VIII-E R2R3MYB proteins encoded by *M. polymorpha* show partial conservation of the motif. Specifically, they lack the [D/E] at the first position of the motif and the [R/K] at the fifth position, while the remaining positions are conserved. The conserved residues include the L in the ninth position and the R in the final position. The study of Zimmermann *et al*. (2004) which delineated the motif, found that these were the only residues which consistently had a strong effect on interaction strength when mutated. A weaker and variable effect was seen when the [R/K] at the fifth position, which is not conserved in the *M. polymorpha* sequences, was mutated. As such, the *M. polymorpha* orthologues of MBW-forming MYB genes exhibit partial conservation of the interaction motif.

**Fig 1.**
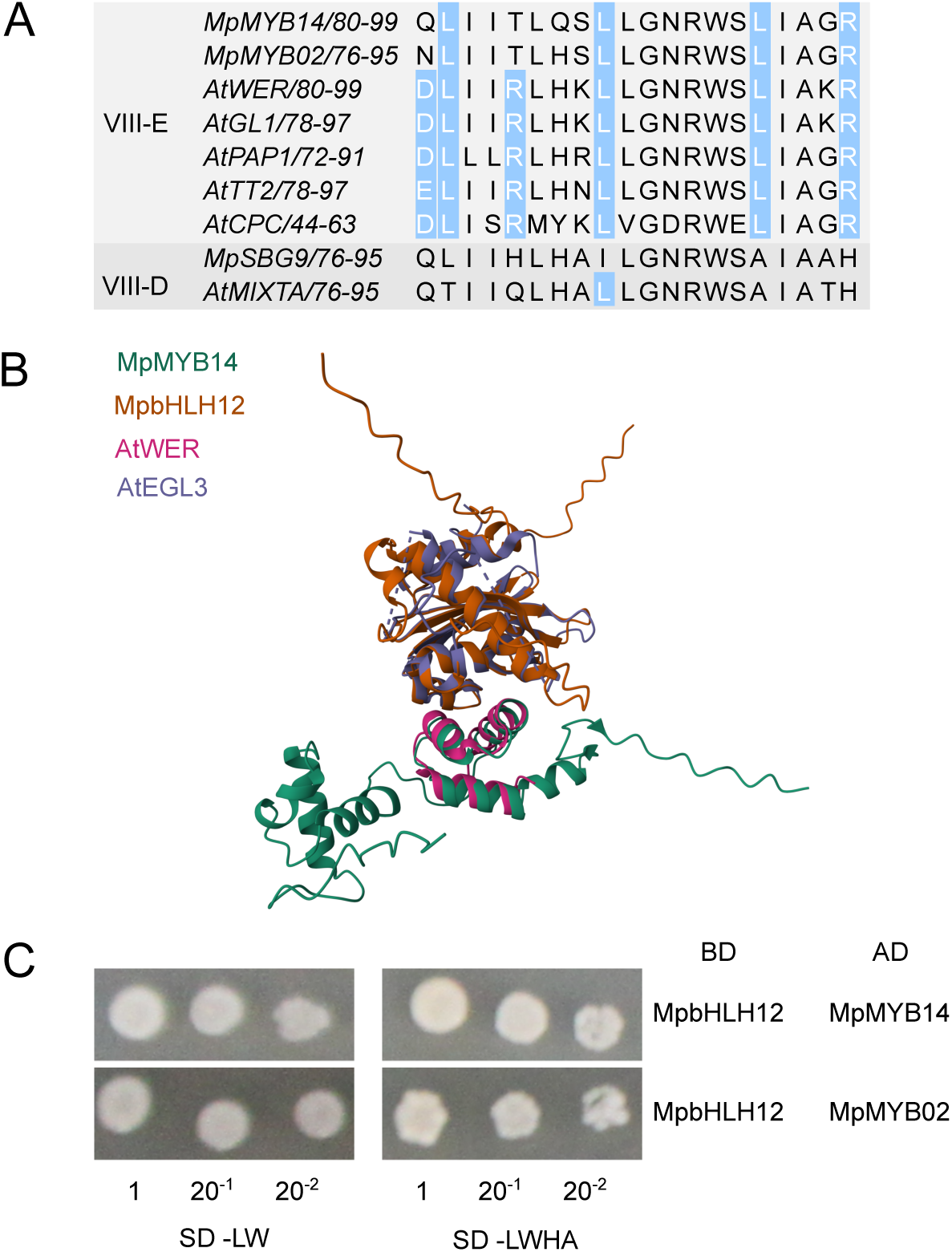
A partially conserved motif can mediate interactions of each of the two VIII-E MYB proteins with the single IIIf bHLH protein of Marchantia polymorpha. **A.** Amino acid sequence alignment of a region of the R3 repeat of MYB proteins from M. polymorpha and A. thaliana. A motif (highlighted in blue) in this region is reported to mediate interaction with subfamily IIIf bHLH proteins in Arabidopsis thaliana (Zimmermann et al., 2004). Four A. thaliana subfamily VIII-E R2R3MYB proteins are shown (AtWER, AtGL1, AtPAP1, AtTT2) and one single-repeat R3MYB protein (AtCPC). The two members of subfamily VIII-E encoded by M. polymorpha exhibit partial conservation of the motif, lacking conservation of the first and fifth residue positions. The motif is restricted to this subfamily, as illustrated by two members of the sister clade VIII-D, from each of the two species. **B.** An AlphaFold Multimer dimer prediction for the MpbHLH12 N-terminal region (aa 1-220) and the R2R3MYB domain (aa 1-144) of MpMYB14 was aligned to the solved crystal structure of a dimer between the N-terminal region of AtEGL3 (aa 1-205) and the R3 MYB repeat of AtWER (aa 67-120) (Wang et al., 2022). The same structural alignment for the predicted MpbHLH12-MpMYB02 dimer and the predicted aligned error plots for the dimer predictions are given in Fig S1. **C.** Yeast two-hybrid tests for interaction between the single M. polymorpha IIIf bHLH protein and each of the two VIII-E MYB proteins. One biological replicate out of four is shown in each instance, the rest are given in Fig S2. For each replicate, two serial dilutions of the eluted cells were performed, as shown. Growth on medium without leucine and tryptophan (-LW) confirms that the yeast clones carry both plasmids with the genes of interests in translational fusion with either the Binding Domain (BD) or Activation Domain (AD) of the yeast transcription factor GAL4 (indicated in the table on the right). Here, in both cases, interaction between the proteins of interest allowed a reconstituted GAL4 to drive expression of reporter genes from the GAL4 Upstream Activation Sequence (UAS). This reporter gene activity complemented two independent auxotrophic mutations in histidine and adenine biosynthesis genes in the background genotype, allowing the yeast to grow in the absence of these amino acids (-HA). Further tests for interaction strength using 3-amino-1,2,4-triazole, a competitive inhibitor of the gene product of the histidine biosynthesis reporter, are shown in Fig S2.

As a first step to investigate whether these motifs may mediate interaction in *M. polymorpha,* we performed *in silico* structural predictions. A predicted interaction interface exhibiting low predicted aligned error was identified between the R3 MYB repeat of both of MpMYB14 and MpMYB02 and the N-terminal region of the single endogenous IIIf bHLH MpbHLH12 (Fig S1A). This recapitulates the experimentally determined interacting regions for the *A. thaliana* complexes (Zhang *et al*., 2003; Wang *et al*., 2022). When an VIII-D MYB, AtMIXTA or MpSBG9, was instead included in the multimer prediction, the predicted aligned error in this region rose markedly (Fig S1B), indicating that an interaction interface was not confidently predicted. In Fig 1B, the predicted conserved interaction can be appreciated in a structural alignment of the predicted MpbHLH12-MpMYB14 dimer on the solved crystal structure of the IIIf bHLH AtEGL3 and VIII-E MYB AtWER. The same visualisation is shown for the predicted MpbHLH12-MpMYB02 dimer in Fig S1A. Together, structural analysis predicts that the interaction interface is conserved for the *M. polymorpha* orthologues, in spite of partial conservation of the sequence motif associated with the homologous interaction in *A. thaliana*.

To validate these predictions *in vivo*, we expressed our proteins of interest in *Saccharomyces cerevisiae AH109* in translational fusion with domains of the *S. cerevisiae* transcription factor GAL4, to perform yeast two-hybrid assays. For both predicted *M. polymorpha* MYB-bHLH dimers, four biological replicates all showed growth indicating sufficient GAL4 reporter activation to complement the strain’s auxotrophy for both histidine and adenine (Fig 1C) and, in the absence of histidine alone, addition of at least 1 mM of a competitive inhibitor of the histidine biosynthesis reporter gene, 3-amino-1,2,4-triazole (Fig S2). This demonstrates that stable protein-protein interactions can form between MpbHLH12 and each of MpMYB14 and MpMYB02 in *S. cerevisiae*.

Having found that these MYB-bHLH complexes could form when expressed heterologously, we sought to determine whether they are co-expressed in their native context. For this purpose, we generated *M. polymorpha* lines carrying transcriptional reporters as described in Romani *et al*. (2024). In the asexual propagules, gemmae, the Mp*MYB02* reporter showed specific expression in oil body cells (Fig 2A), as has recently been reported independently (Wang *et al*., 2023) and is consistent with the reported role of Mp*MYB02* in the maturation of the oil body. Signal from the Mp*MYB14* reporter was strongest around the notch regions in dormant gemmae and later became more widely distributed across the thallus (Fig 2B), consistent with a published GUS reporter line (Albert *et al*., 2018). Notably, we did not observe signal from the Mp*MYB14* reporter in the oil body cells, indicating that the expression patterns of these two MYB paralogues are mutually exclusive. The Mp*bHLH12* reporter showed strong expression in a broad domain which was inclusive of both of the more specific *MYB* expression domains (Fig 2C). Based on these observations, it is possible that MpbHLH12 could dimerise with each of MpMYB14 and MpMYB02 in each of their distinct expression domains.

**Fig 2.**
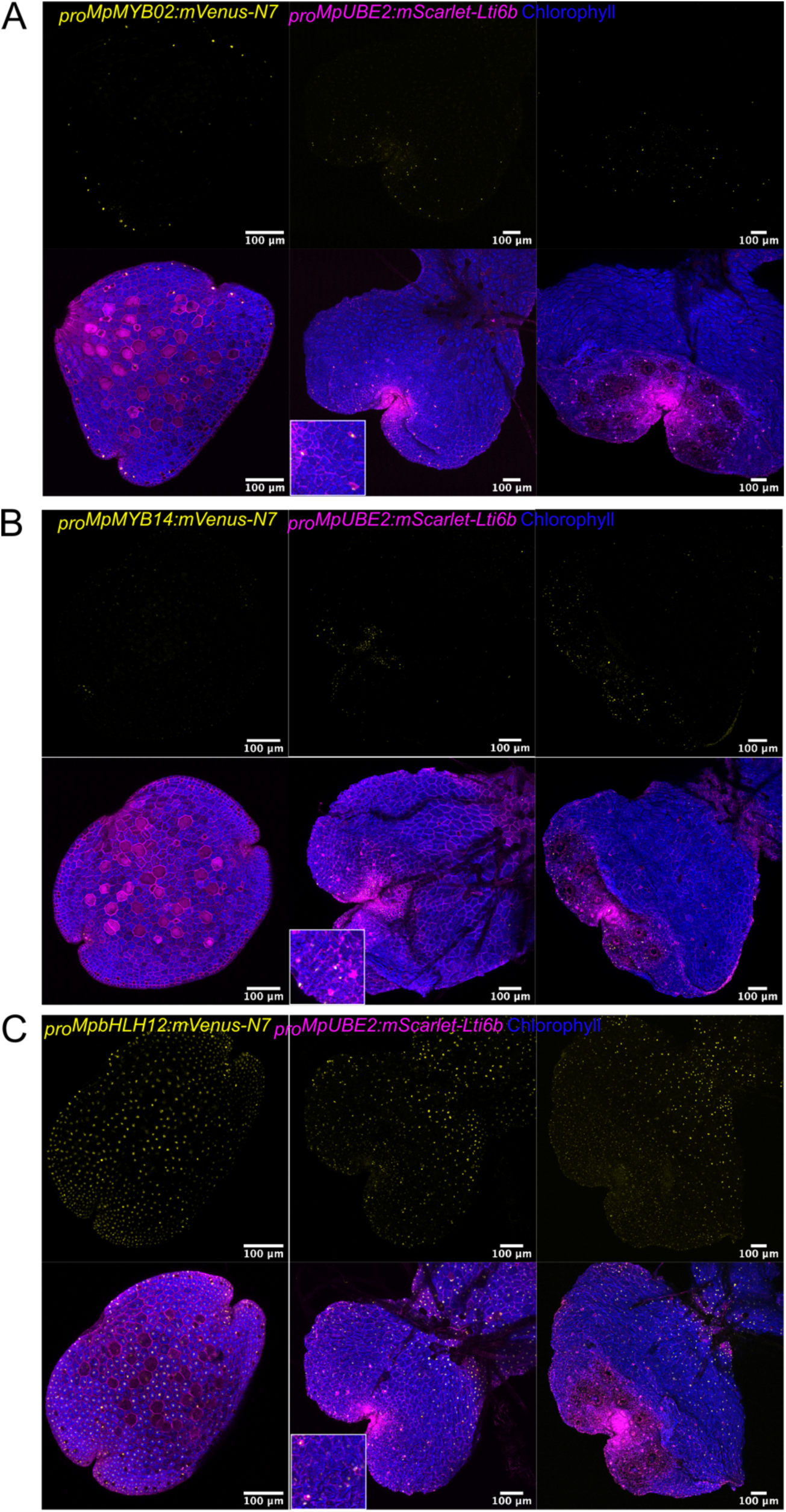
A transcriptional reporter for the single Marchantia polymorpha IIIf bHLH gene is broadly expressed across developmental stages, encompassing both of the specific regions where each of the two VIII-E MYB genes are expressed. Confocal micrograph time series of transcriptional reporter lines for MpMYB02 (**A**), MpMYB14 (**B**) and MpbHLH12 (**C**). Gemmae were analysed of isogenic lines deriving from Cam1x2 spores with two independent transgenes integrated, an mScarlet plasma membrane marker (shown in magenta) and the proximate 5’ cis-regulatory region of each of the genes of interest driving the expression of nuclear-localised mVenus (shown in yellow). Chlorophyll autofluorescence was also captured in each case (shown in blue). Maximum intensity projections are shown of z-stacks for the mVenus channel only alongside merged images from all three channels. Imaging was done on days 0, 3 and 5 after gemma germination. Insets show higher magnification of oil body cells.

### The functions of the two *Marchantia polymorpha VIII-E MYB* genes both depend on the single *IIIf bHLH* gene

Previous studies have demonstrated the requirement for Mp*MYB14* and Mp*MYB02* for auronidin pigment production and oil body maturation, respectively. To test our hypothesis that each of the encoded MYB proteins form transcriptional complexes with the single *M. polymorpha* IIIf bHLH to carry out their functions, we generated gene knockout lines using CRISPR/Cas9 (Fig S3) and scored these phenotypes. Consistent with reports from other accessions (Albert *et al*., 2018; Carella *et al*., 2019), we could induce strong auronidin pigmentation in plants deriving from a cross between the *Cam-1* and *Cam-2* accessions by transfer of four-week old plants to minimal medium (Fig 3A), and we found that this response was abolished in several independent Mp*myb14* loss-of-function mutant lines (Fig 3A and S4). The response was not affected in loss-of-function mutants for the paralogous Mp*myb02* (Fig 3A and S4). However, the response was abolished in four independent Mp*bhlh12* loss-of-function mutant lines (Fig 3A and S4). Auronidin production could be rescued in an Mp*bhlh12* line by introduction of Mp*bHLH12* (Fig 3A and S4). In conclusion, Mp*bHLH12* is required for the Mp*MYB14*-activated auronidin pigment response, while Mp*MYB02* is not involved.

**Fig 3.**
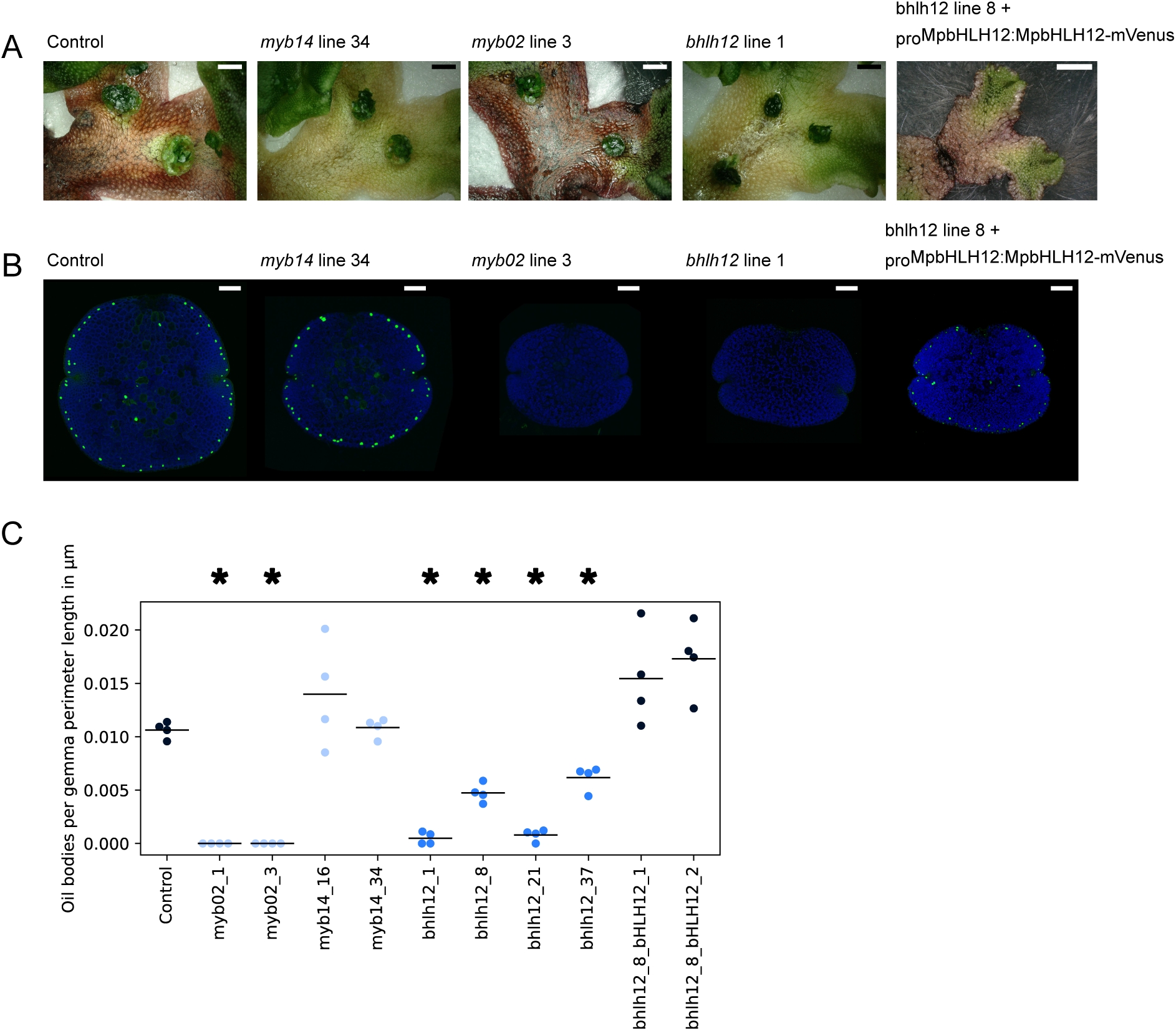
Loss-of-function of the single IIIf bHLH gene causes both the lack of auronidin pigment and the impaired oil body maturation seen in each of the loss-of-function mutants for the two VIII-E MYB genes in Marchantia polymorpha. Phenotypes of representative loss-of-function CRISPR/Cas9 mutants at each of the three loci. An isogenic Cam1x2-derived line bearing a MpTOC1:LUC reporter was used as a non-edited transgenic control line. **A**. Auronidin pigmentation was scored in lines grown under nutrient deprivation (see Methods). Three additional independent CRISPR/Cas9 Mpbhlh12 knockout lines showing the same phenotype and a line with an in-frame mutation in MpbHLH12 showing the wild-type phenotype are shown in Fig S4. An additional independent line for each of Mpmyb14 and Mpmyb02 and an additional Mpbhlh12 MpbHLH12 complementation line showing the same phenotypes are also shown in Fig S4. Scale bars are 2 mm. **B.** Oil bodies were counted in dormant gemmae stained with the fluorescent lipid stain BODIPY and imaged with a confocal microscope. One gemma is shown for each genotype. Scale bars are 100 μm. **C.** The number of mature oil bodies was scored in four BODIPY-stained gemmae for each line. For each gemma, the count was normalised by the perimeter length, to account for differences in gemma size. The raw counts and perimeter measurements are given in the Supplemental Materials. Each line was compared to the control line using a rank-based one-way analysis of variance, the Kruskal-Wallis test, and those significantly reduced in oil body counts per perimeter length at p < 0.05 are marked with an asterisk. The points are coloured according to their genotypes, with control in dark blue, bHLH mutants in medium blue, and MYB mutants in light blue.

To assess the maturation of oil bodies in these lines, we stained dormant gemmae with the fluorescent lipid stain BODIPY and imaged them using confocal microscopy. No BODIPY-staining oil bodies were found in our Mp*myb02* mutant lines (Fig 3B and C), consistent with the reported lack of mature oil bodies in Mp*myb02* loss-of-function mutants (Wang *et al*., 2023). Mutants for the paralogous Mp*MYB14* did not differ from the control (Fig 3B and C). However, Mp*bhlh12* mutants exhibited a strong reduction in the number of mature oil bodies. The severity of the phenotype varied within and between lines (Fig 3B and C). Again, we confirmed that complementation with Mp*bHLH12* could rescue the number of BODIPY-positive oil bodies in an Mp*bhlh12* mutant line (Fig 3B and C). In conclusion, Mp*bHLH12* also aids the Mp*MYB02*-dependent maturation of oil bodies, although in this case requirement was partial. Additionally, we conclude that the mutant phenotypes of the two *VIII-E MYB* genes are distinct. Overall, this is consistent with the hypothesis that MpbHLH12 forms two alternative complexes with MpMYB14 and MpMYB02 in each of the distinct expression domains of the two *MYB* genes.

### Two *Marchantia polymorpha* TTG1-like WDR proteins can form complexes with the IIIf bHLH protein

Having found evidence indicating that complexes form between the single IIIf bHLH and the two VIII-E MYB proteins of *M. polymorpha*, we asked whether these are scaffolded by TTG1-like WDR proteins, as the homologous complexes are in seed plants. As a first step in testing this, we performed structural predictions of trimers. For all of the TTG1 homologues we tested from *A. thaliana* (AtTTG1 and AtLWD1) and *M. polymorpha* (MpWDR1, MpWDR2 and MpWDR3), a region of low predicted aligned error was found, encompassing a region just N-terminal to the bHLH domain of the bHLH protein and the entire highly conserved beta propeller structure of the WD repeat protein (Fig S1B). This region is highlighted in the predicted structures for AtTTG1-AtTT8-AtTT2 and MpWDR1-MpbHLH12-MpMYB14 in Fig 4A. Note that in *A. thaliana*, AtTTG1 on its own is required for all MBW complex activities and its paralogues AtLWD1 and 2 cannot complement *ttg1* mutants when ubiquitously expressed (Airoldi *et al*., 2019). Using yeast two-hybrid assays, we found that MpWDR1 and 2, but not 3, interacted with MpbHLH12 (Fig 4B). As such, AlphaFold predicted that all of the TTG1 paralogues and orthologues tested have the capacity to scaffold MBW complexes, but *in vivo* evidence indicates that only some paralogues do in each lineage, AtTTG1 in *A. thaliana* (Airoldi *et al*., 2019) and MpWDR1 and 2 in *M. polymorpha* (Fig 4B).

**Fig 4.**
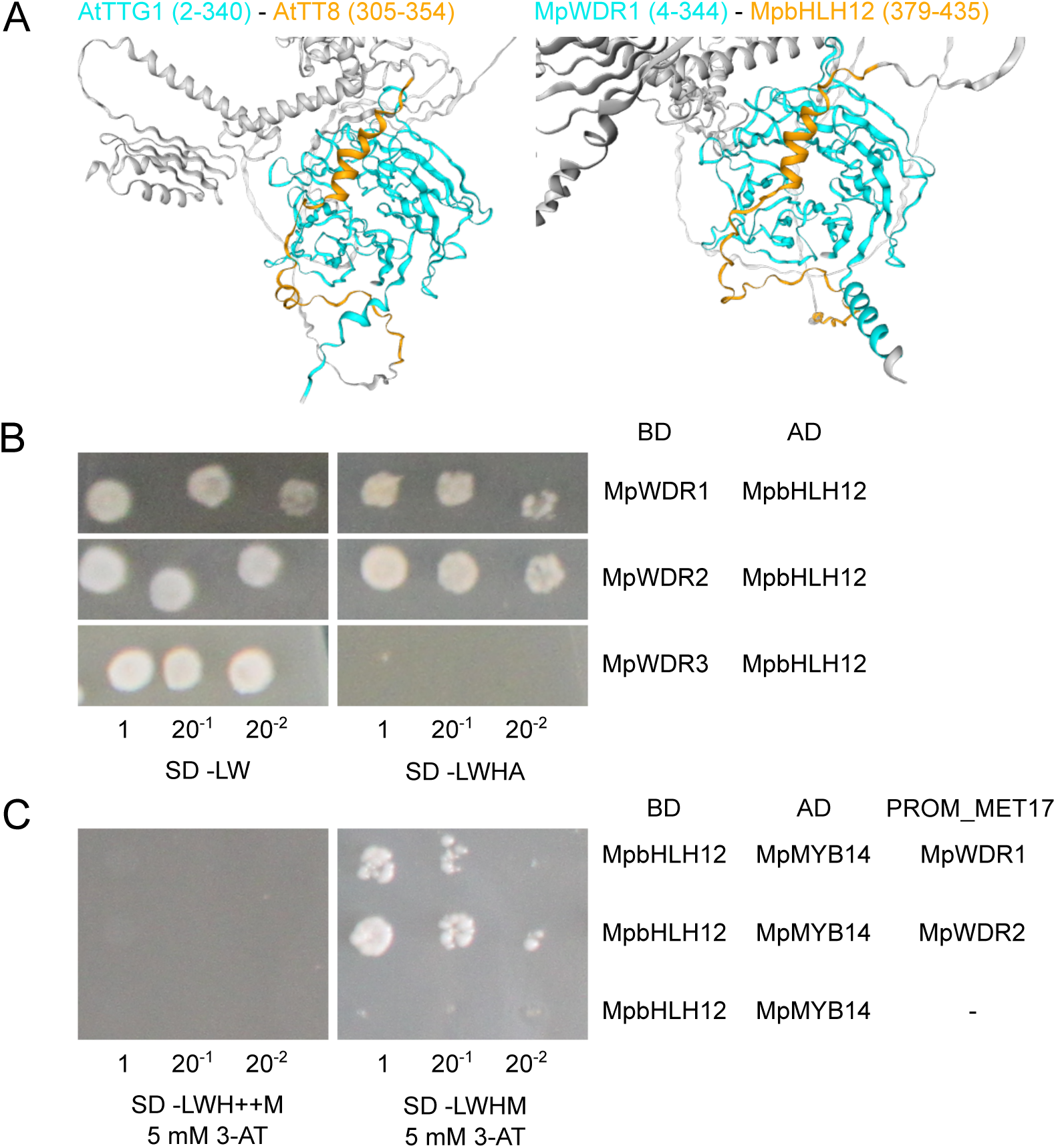
Marchantia polymorpha TTG1-like WDR proteins interact with the single IIIf bHLH protein and may stabilise tripartite MBW complexes. **A.** AlphaFold Multimer trimer predictions indicate that TTG1-like WDR proteins from M. polymorpha can engage in tripartite complexes with the idenfitied MYB-bHLH dimers through interaction of the conserved WDR beta-propeller with a region just N-terminal to the bHLH domain. This predicted interaction interface, which exhibits a low predicted aligned error score (full plots given in Fig S1) is highlighted. **B.** Yeast two-hybrid tests for interaction between the single M. polymorpha IIIf bHLH and each of the three M. polymorpha TTG1-like WDR proteins. One biological replicate out of four is shown in each instance, the rest are given in Fig S2. For each replicate, two serial dilutions of the eluted cells were performed, as shown. Growth on medium without leucine and tryptophan (-LW) confirms that the yeast clones carry both plasmids with the genes of interests in translational fusion with either the Binding Domain (BD) or Activation Domain (AD) of the yeast transcription factor GAL4 (indicated in the table on the right). Here, for each of MpWDR1 and 2 paired with MpbHLH12, interaction between the proteins of interest allowed a reconstituted GAL4 to drive expression of reporter genes from the GAL4 Upstream Activation Sequence (UAS). This reporter gene activity complemented two independent auxotrophic mutations in histidine and adenine biosynthesis genes in the background genotype, allowing the yeast to grow in the absence of these amino acids (-HA). Further tests for interaction strength using 3-amino-1,2,4-triazole, a competitive inhibitor of the gene product of the histidine biosynthesis reporter, are shown in Fig S2. No interaction sufficient to drive reporter gene activation allowing growth in these conditions was observed between MpWDR3 and MpbHLH12. **C.** Conditional expression assay to test the effect of presence of a WDR protein on the MYB-bHLH interaction strength. One biological replicate out of four is shown in each instance, the rest are given in Fig S2. In the presence of methionine (++M), the MET17 promoter is inactive and growth of the MpbHLH12-MpMYB14 two-hybrid on -LWH could be abolished with the addition of 5 mM of 3-AT. In the absence of methionine (-M), the MET17 promoter is active and drives the expression of one of the WDR genes. For MpWDR1 and 2, this allowed growth at ≥5 mM 3-AT (although variation was observed between biological replicates for MpWDR1, see Fig S2), while the absence of methionine had no effect on the growth of the two-hybrid without the MET17 cassette.

To test whether the *M. polymorpha* orthologues that interact in pairwise MYB-bHLH and bHLH-WDR dimers form tripartite MYB-bHLH-WDR complexes, we modified our yeast assay to allow us to interrogate the effect of inducible expression of a third protein on the strength of interaction between two others. Noisiness of growth under the highly restrictive conditions in this assay prevented reliable interpretation for some combinations of proteins tested, but we identified signal indicating that the presence of either MpWDR1 or 2 enhances the MpbHLH12-MpMYB14 interaction (Fig 4C). This was consistent across all four biological replicates for MpWDR2, but varied between replicates for MpWDR1 (Fig S2). This is indicative of cooperative dynamics, as reported for the AtTT2-AtTT8-AtTTG1 MBW complex (Baudry *et al*., 2004).

We also generated transcriptional reporter lines for Mp*WDR1* and *2*. The Mp*WDR1* reporter was strongly and ubiquitously active across the stages observed, including in oil body cells (Fig 5A). The Mp*WDR2* reporter was broadly expressed but dim (Fig 5B). These observations are consistent with RNA-seq experiments across vegetative tissues (Kawamura *et al*., 2022). This indicates that expression of both of these proteins overlaps with those of the two MYB-bHLH dimers and that trimers could form *in planta*.

**Fig 5.**
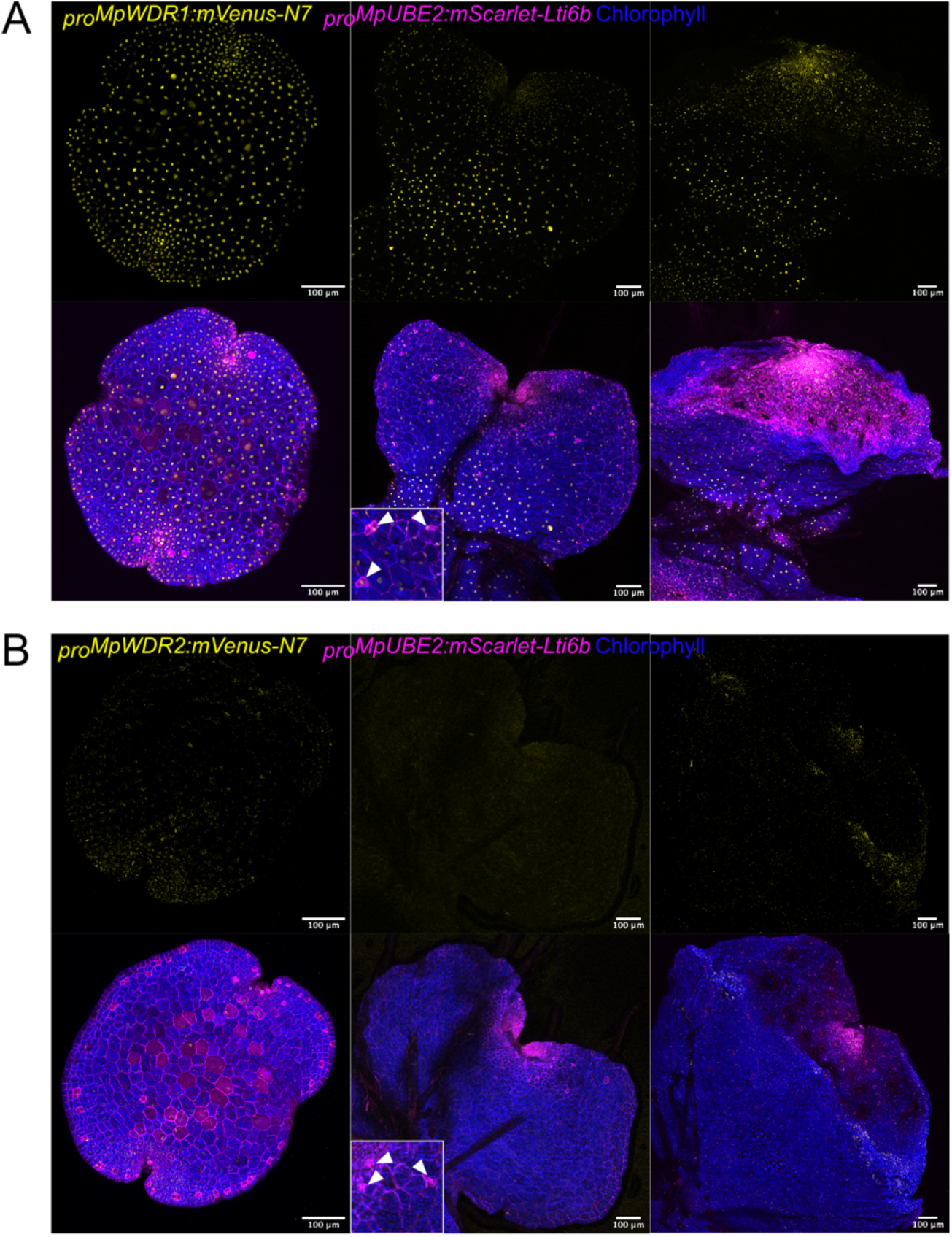
Transcriptional reporters for the two Marchantia polymorpha TTG1-like WDR genes whose protein products interact with the MYB-bHLH dimers drive broad expression, but their levels differ. Confocal micrograph time series of transcriptional reporter lines for MpWDR1 (**A**) and MpWDR2 (**B**). Gemmae were analysed of isogenic lines deriving from Cam1x2 spores with two independent transgenes integrated, an mScarlet plasma membrane marker (shown in magenta) and the proximate 5’ cis-regulatory elements of each of the genes of interest driving the expression of nuclear-localised mVenus (shown in yellow). Chlorophyll autofluorescence was also captured in each case (shown in blue). Maximum intensity projections are shown of z-stacks for the mVenus channel only alongside merged images from all three channels. Imaging was done on days 0, 3 and 5 after gemma germination. Insets show higher magnification of oil body cells.

### Low recovery of loss-of-function alleles prevented the investigation of higher-order mutants for the *Marchantia polymorpha TTG1-like WDR* genes

To test whether TTG1-like WDR scaffolds are required for the functions of the identified MYB-bHLH dimers in *M. polymorpha*, we set out to generate loss-of-function mutants by targeting the loci of the three paralogues with CRISPR/Cas9 (Fig S3). Given the data presented here and that of Airoldi *et al*. (2019), we reasoned that Mp*WDR1* and 2 may have a redundant role in scaffolding MBW complexes, while the third paralogue, Mp*WDR3*, is likely divergent in function. Based on this, we devised strategies to target the Mp*WDR1* and *2* loci simultaneously, all three loci simultaneously and only Mp*WDR3* (see Methods). We recovered predicted loss-of-function single mutants for Mp*WDR2* and Mp*WDR3*, but not for Mp*WDR1*. Among 80 primary transformants genotyped at that locus, two edited Mp*WDR1* alleles were identified. One harboured a 3 bp deletion which likely did not affect protein function. Another exhibited a 1 bp deletion causing an early frameshift. This putative loss-of-function allele was identified in a line in which a predicted loss-of-function mutation was also detected at Mp*WDR2*. This Mp*wdr1,2* double mutant line was the only higher-order mutant line recovered.

The single mutants, Mp*wdr2* and Mp*wdr3*, exhibited no marked impairment for the traits regulated by the MYB-bHLH dimers (Fig 6). A mild, statistically significant reduction in the number of mature oil bodies in gemmae was observed in only one of the two independent *wdr3* mutant lines (Fig. 6B). The double Mp*wdr1,2* mutant line showed severe growth defects making it difficult to compare its phenotypes to other lines, but auronidin production was functional and some putative BODIPY-staining oil bodies were observed (Fig 6). Note that for this genotype only a single line was recovered, so the effects of these mutations could not be evaluated in an independent genomic background.

**Fig 6.**
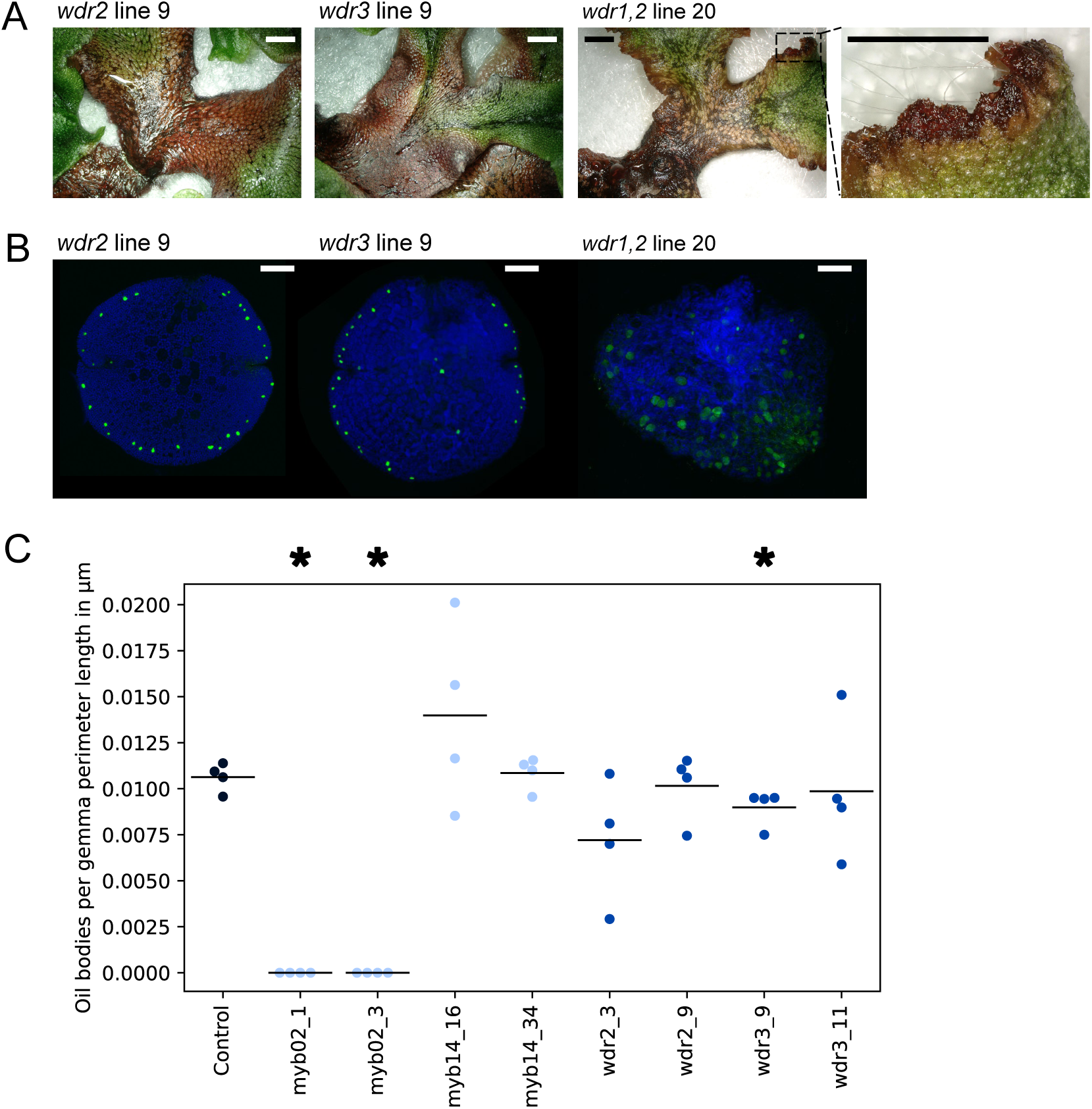
Loss-of-function mutant lines could only be recovered for some of the Marchantia polymorpha TTG1-like WDR genes, and those produce auronidins and show no consistent impairment in oil body maturation. Phenotypes of representative loss-of-function CRISPR/Cas9 mutants at the WDR loci. An isogenic Cam1x2-derived line bearing a MpTOC1:LUC reporter was used as a non-edited transgenic control line. **A.** Auronidin pigmentation was scored in lines grown under nutrient deprivation (see Methods). An additional independent CRISPR/Cas9 knockout line for each gene showing the same phenotype is given in Fig S4 for the single mutants, while for the double mutant, only a single line was recovered. Scale bars are 2 mm. **B.** Oil bodies were counted in dormant gemmae stained with the fluorescent lipid stain BODIPY and imaged with a confocal microscope. One gemma is shown for each genotype. Scale bars are 100 μm. **C.** The number of mature oil bodies was scored in four BODIPY-stained gemmae for each line. For each gemma, the count was normalised by the perimeter length, to account for differences in gemma size. The raw counts and perimeter measurements are given in the Supplemental Materials. Each line was compared to the control line using a rank-based one-way analysis of variance, the Kruskal-Wallis test, and those significantly reduced in oil body counts per perimeter length at p < 0.05 are marked with an asterisk. Note that the control and MYB mutant samples are the same as displayed in Fig 3. The points are coloured according to their genotypes, with control in dark blue, WDR mutants in medium blue, and MYB mutants in light blue.

In conclusion, we found that neither of Mp*WDR2* or Mp*WDR3* were singly required for the activities of the MYB-bHLH complexes. For the final paralogue, the highly expressed Mp*WDR1*, no loss-of-function single mutants were recovered, indicating that mutations at this locus have pleiotropic effects on viability. No triple mutant lines and only a single double mutant line were recovered. This Mp*wdr1*,*2* line exhibited growth defects, and could not be replicated, indicating that more severe defective phenotypes may be suppressed in this line. As such, we were not able to evaluate the individual effect of mutation of Mp*WDR1*, nor to address whether there is functional redundancy between the three paralogues. In conclusion, we could not use functional data to confirm whether TTG1-like WDR scaffold proteins are required for the activities of the identified MYB-bHLH complexes in *M. polymorpha*.

## Discussion

### A MYB-bHLH transcriptional complex is ancestral to the land plants

MBW transcriptional complexes regulate a diverse set of pathways in the elaboration of the epidermis of seed plants (Ramsay and Glover, 2005; Nemesio-Gorriz *et al*., 2017). The MYB member of the complex has been highlighted as a “hotspot” locus underlying the evolution of pigmentation phenotypes across angiosperms (Sobel and Streisfeld, 2013; Wheeler *et al*., 2022; Marin-Recinos and Pucker, 2024). Here we investigated the origin of the complex and its evolution beyond this clade. It has been known for some time that orthologous genes are also encoded by bryophytes, but incomplete conservation of the canonical interaction motif in the R3 MYB repeat (Grotewold *et al*., 2000; Zimmermann *et al*., 2004) had been taken to imply that these do not form MBW complexes (Davies *et al*., 2019). Contrary to this, here we found evidence *in silico* and in yeast that both of the two *M. polymorpha* MYB proteins orthologous to those that form MBW complexes in seed plants, MpMYB14 and MpMYB02, interact with the single endogenous IIIf bHLH, MpbHLH12. Further, we generated CRISPR/Cas9 loss-of-function mutants for this *bHLH* gene and found that these lines phenocopied both of the distinct phenotypes of mutants for each of the *MYB* genes. This suggests that the formation of these complexes is required, partially in the case of oil body maturation and wholly in the case of auronidin pigmentation, for the functions of the MYB proteins. In another member of the order Marchantiales, *Plagiochasma appendiculatum*, a single IIIf bHLH has also been identified that is associated both with the production of flavonoids and bis-bibenzyls, the latter of which accumulate in oil bodies (Wu *et al*., 2018). In *M. polymorpha*, we found that the *bHLH* was broadly expressed, encompassing both of the non-overlapping expression domains of the *MYB* genes. As such, we propose that two distinct MYB-bHLH transcriptional complexes form in each of these domains in *M. polymorpha*. Taking our new data into account, we can trace back the origin of the MYB-bHLH interaction all the way to the origin of the *VIII-E MYB* genes and the *IIIf bHLH* genes, in the stem lineage of land plants (Jiang and Rao, 2020; Kongsted and Glover, 2023).

As for the third member of the complex, the WDR scaffold protein, we found evidence in yeast expression systems that two of the TTG1-like WDR proteins in *M. polymorpha*, MpWDR1 and 2, but not the third paralogue MpWDR3, can engage in complexes with these MYB-bHLH dimers. Airoldi *et al*. (2019) found that the same paralogues, MpWDR1 and 2, but not 3, could complement *ttg1* mutant phenotypes when heterologously expressed in *Arabidopsis thaliana*. Taken together, these analyses suggest that the same features are involved in interaction of TTG1-like WDR proteins with the native IIIf bHLH in *M. polymorpha* as with heterologous partners in *A. thaliana*. This suggests that a TTG1-like WDR protein acted as a scaffold for the MYB-bHLH complexes ancestrally. Using several CRISPR/Cas9 approaches, we found that both Mp*WDR2* and *3* individually are dispensable for the functions of the two MYB-bHLH complexes. However, due to suspected effects on viability of mutation of Mp*WDR1*, we were unable to recover higher-order mutant lines to determine whether this is due to redundancy between the *TTG1-like WDR* genes in *M. polymorpha*. In *A. thaliana*, all of the TTG1-like WDR genes have additional functions unrelated to MBW complexes and the triple mutant (*ttg1 lwd1 lwd2*), in addition to a loss of pigments and mispatterning of epidermal cell types, shows complete arrhythmicity of the circadian clock, late flowering and an aberrant leaf morphology (Airoldi *et al*., 2019). It seems likely that similar pleiotropic effects hindered the recovery of mutants in *M. polymorpha*.

In conclusion, we propose that the best-supported hypothesis is that the MBW complex originated in the stem lineage of land plants. Other examples have been identified of transcriptional complexes conserved across similar timescales of plant evolution, such as the BONOBO-LRL bHLH heterodimer that is an ancestral specifier of land plant germ cells (Saito *et al*., 2023) and the Ia-IIIb bHLH heterodimer that regulates stomatal development across land plants (Chater *et al*., 2016; Harris *et al*., 2020). Even more ancient is the reported conserved KNOX-BELL TALE homeodomain heterodimer involved in development in the diploid generation across the Viridiplantae (Dierschke *et al*., 2021). However, in all of these cases the complexes are composed of members of the same protein family and could thus conceivably have evolved from homomeric interactions. To our knowledge, the present report is the first example that involves members of multiple different protein families, highlighting the flexible evolution of regulatory interactions between distantly related proteins in plants.

### Homologous regulation of convergent flavonoid pigments

Our work demonstrates that homologous MYB-bHLH transcriptional complexes regulate the flavonoid pathway in *M. polymorpha* and in the seed plants. Red pigments are produced from this pathway in all major lineages of land plants, with the exception of the hornworts (Davies *et al*., 2022). We could not detect any instances of the LX_6_LX_6_LX_3_R MYB R3 interaction motif in the genome of the hornwort *Anthoceros agrestis* (Li *et al*., 2020) and *A. agrestis* also lacks the IIIf subclass of bHLH proteins (Kongsted and Glover, 2023). Together, our data are consistent with a single ancestral origin of MBW regulation of the flavonoid pathway in the land plants, and the absence of these regulators in the hornworts is associated with the absence of red pigmentation. However, the red flavonoid pigments produced across lineages are not homologous. While the early steps of the flavonoid biosynthetic pathways are shared, distinct later steps lead to the production of 3-hydroxyanthocyanins in seed plants, 3-deoxyanthocyanins in ferns and mosses, and auronidins in liverworts (Davies *et al*., 2022). As such, the ancestral condition is not yet clear, but, as has been posited elsewhere (Bowman, 2022), it appears likely that a red pigment was produced from the flavonoid pathway in stem land plants. Notably, Bowman (2022) also speculated that this hypothetical ancestral pigment was regulated by a MYB transcription factor. Regardless of the ancestral red pigmentation pathway (if any), all the steps of the flavonoid biosynthetic pathway required for the production of UV-screening flavones have been inferred to have been present in the last common ancestor of the land plants (Davies *et al*., 2022). Our findings support the hypothesis that these, and perhaps other products of the flavonoid pathway present in the common ancestors of extant land plants, were regulated by an MBW transcriptional complex.

### Lineage-specific complexes have evolved by an independently repeated mechanism

In addition to the flavonoid pathway, MBW complexes which regulate lineage-specific traits have been identified in multiple taxa. For instance, in the rosid lineage of angiosperms, MBW complexes regulate the patterning of unicellular trichomes on aerial organs (Serna and Martin, 2006). Here, we find that homologous complexes in liverworts are involved in the maturation of oil bodies, a synapomorphy of the liverwort lineage. In both cases, the gain of a new function is associated with a lineage-specific duplication of the *MYB* member and co-option of its interaction partners. In the angiosperms, a duplication gave rise to *VIII-E MYB* subgroups 6 vs 15 (Jiang and Rao, 2020) and the bHLH AtEGL3 regulates both anthocyanin biosynthesis and trichome development by interaction with MYB proteins from either subgroup (Zhang *et al*., 2019). The Mp*MYB14* vs Mp*MYB02* duplication appears to be liverwort-specific (Jiang and Rao, 2020) and both form a complex with MpbHLH12, which regulates both their functions. As a result of this pattern observed across lineages, the MYB members remain pathway-specific, while the other members act across multiple pathways. This pleiotropy may constrain their evolution and explain the observed bias towards changes to the *MYB* member of the complex (Sobel and Streisfeld, 2013; Wheeler *et al*., 2022; Marin-Recinos and Pucker, 2024). Other examples have been reported of MBW orthologues that may have lineage-specific functions in a number of angiosperms. For example, specific expression of *VIII-E MYB* genes has been identified in idioblasts of bright eyes (*Catharanthus roseus*) (Li *et al*., 2024) and *Citrus* oil glands (Wang *et al*., 2024). Interestingly, trichomes, oil glands, and oil bodies play analogous roles as secretory structures which allow the accumulation of toxic compounds to defend against herbivory, but they have evolved independently in each lineage (Romani *et al*., 2022). Further, *MYB* genes with complete conservation of the canonical interaction motif within the R3 MYB repeat are found across land plants, notably in mosses (in the genome of *Physcomitrium patens*) and in non-seed tracheophytes (in the genome of *Selaginella moellendorffii*) (Jiang and Rao, 2020). These species also encode *IIIf bHLH* genes (Kongsted and Glover, 2023) and *TTG1-like WDR* genes (Airoldi *et al*., 2019). As such, there is ample opportunity across phylogenetic contexts to test whether the observed patterns in the evolution of these transcriptional complexes are independently replicated.

## Supporting information

Raw counts

## Acknowledgements

We are grateful to Eva Herrero Serrano for help with establishing protocols for protein-protein interaction tests in the lab and for discussions about the evolution of protein functions. We thank Susana Sauret-Güeto, Jenna Rever, Davide Annese and David Hoey for additional help with *M. polymorpha* protocols. For discussion of the project as it developed, we want to thank Nathanael Walker-Hale, Sam Brockington, Alex Webb, Hanna Marie Schilbert, Julia Davies and Sebastien Andreuzza. We are grateful to Matthew Dorling and Qi Wang for support in the lab and on the computing cluster, respectively. This work was funded by UKRI Natural Environment Research Council doctoral training programme studentship 2277410 to TEK, Biotechnology and Biological Sciences Research Council grant BB/T007117/1 to JH and BBSRC/EPSRC OpenPlant Synthetic Biology Research Centre BB/L014130/1 to JH. FR is a Leverhulme Early Career Fellow.

## Author contributions

T.E.K. lead the project, performed the majority of the experiments and analysis, interpreted the data and wrote the paper with input from all authors. F.R. performed the transcriptional reporter experiments and the complementation experiments and helped with genetic transformation of *M. polymorpha*. C.A.A. cloned genes for expression in yeast, performed preliminary experiments and provided training and supervision to T.E.K. J.H. shared protocols and resources and acquired funding. B.J.G. supervised the project and acquired funding.

