## Supplemental materials for "Replicated repurposing of an ancestral transcriptional complex in land plants"

### Materials and methods

#### Plant materials and growth conditions

*Marchantia polymorpha ruderalis* plants deriving from crosses of the Cambridge isolates, *Cam-1* (male) and *Cam-2* (female) were maintained in a growth room in long-day conditions at 22-23°C, except where otherwise stated.

#### Analysis of protein sequences and structures

MYB protein sequences were aligned with MAFFT (Katoh and Standley, 2013) and visualised with Jalview (Waterhouse *et al.*, 2009). Structural predictions of protein multimers were generated with AlphaFold2 (Evans *et al.*, 2021) using Colabfold (Mirdita *et al.*, 2022) on the CLIP CBE cluster (https://www.clip.science) at the Vienna Biocenter. Predicted aligned error plots were generated with PAE viewer (Elfmann and Stülke, 2023). Structural alignments were inferred using the RCSB Protein Data bank tool (Bittrich *et al.*, 2024) with default settings.

#### Cloning and assembly for expression in yeast

Mp*bHLH12* (Mp2g05070), Mp*MYB14* (Mp5g19050), Mp*MYB02* (Mp3g07510), Mp*WDR1* (Mp5g03900), Mp*WDR2* (Mp5g01830) and Mp*WDR3* (Mp5g06230) were amplified from *M. polymorpha* cDNA prepared by RNA extraction with TRIzol (Invitrogen), DNAse I treatment and reverse transcription with Bioline Reverse Transcriptase. The cDNA was used as a template for Phusion HF two-step PCR with primers AC11-14, 39-40, 72-75, 191 and 192. These PCR reactions were then used as the template for Phusion HF two-step PCR reactions with the primers AC87-88 to add attB sites. Amplicons were gel purified with the Invitrogen PureLink Quick Gel Extraction kit and integrated into the Gateway entry vector pDonr201 with the Invitrogen Gateway BP Clonase II Enzyme Mix. Presence of the desired insert was confirmed by Sanger sequencing with attL1 and attL2. LR reactions for integration into the destination vectors pAS2 and pC-ACT2 were carried out with the Invitrogen Gateway LR Clonase II Enzyme Mix. To allow inducible expression of a third protein, pAS2 constructs were modified as follows. Phusion HF two-step PCR was used to amplify Mp*WDR1*, *2* and *3* from the pDonr201 entry clones (primers TK9-14), and the methionine-repressible *MET17* promoter (Solow, Sengbusch and Laird, 2005) and the *CYC1* terminator from p406 (primers TK15-18). pAS2-Mp*bHLH12* was digested with AflII and MluI-HF and purified with the Invitrogen PureLink Quick Gel Extraction kit. Parts were integrated into the digested vector by Gibson assembly with the 2X HiFi Assembly Master Mix (NEB). Plasmids were verified by restriction enzyme digestion and by Sanger sequencing with the primers TK50 and M13F. A map of the construct for conditional expression of MpWDR1 is given in Fig S5, as an example. Primer sequences are given in Table S2.

**Yeast transformation and spot testing**

All yeast growth media were prepared according to the Yeast Protocols Handbook (2009, Clontech Laboratories Inc.) except as stated. *Saccharomyces cerevisiae* strain *AH109* was co-transformed with two plasmids, a pAS2 destination plasmid and a pC-ACT2 destination plasmid, by Small-Scale LiAC Yeast Transformation (as in the Yeast Protocols Handbook (2009, Clontech Laboratories Inc.) except YPDA with higher adenine (40 mg/L) was used for the overnight cultures, and in step 8 pellets were resuspended directly in TE/LiAC). For the autoactivation tests, a control plasmid (either the empty destination vector or one with *mVenus* in place of a gene of interest, as indicated in each case) was added instead of the second plasmid. Transformants were selected by growth on -LW dropout medium at 30°C for 5-6 days. When colonies had grown to a size of ~1-2 cm, spot tests were performed as follows. Four colonies were picked from each transformation. Each colony was inoculated in 200 µL sterile 1X TE in wells of a sterile microtitre plate. 10 µL was transferred from inoculated wells to clean wells of 200 µL sterile TE and mixed, and 10 µL was transferred from each of these wells to clean wells of 200 µL sterile TE, to obtain two serially diluted replicates for each colony. 3 µL was plated from each well onto the amino acid dropout media. The media used were -LW, - LWH, - LWHA, and -LWH and where applies were supplemented with 3-amino-1,2,4-triazole (3-AT) for the desired concentration. Spot test plates were grown at 30°C for 4-5 days and imaged with a Canon EOS 500D. As our lab-maintained *AH109* strain was found to require methionine for growth, a fresh stock of Takara Matchmaker *Y2HGold* was maintained on modified -M medium (with 40 mg/L adenine and 100 mg/L aspartic acid and pH adjusted to 5.8), for use with the methionine-repressible expression system. For this, co-transformation was carried out as above, with the following exceptions. In preparation for transformation, the strain was grown on the modified -M medium for ~7 days. ~10-18 colonies (each ~1-2 mm in diameter) were inoculated in liquid modified -M medium and grown at 30°C and 180 rpm for ~24 hours. Transformants were selected on -LWM (with 40 mg/L adenine and 100 mg/L aspartic acid and pH adjusted to 5.8). Four colonies were picked from these plates, streaked out individually on new plates with the same selective medium and grown for six days before spot tests. The media used for spot tests in this case were -LWHM (with 40 mg/L adenine and 100 mg/L aspartic acid), and -LWH++M (with 40 mg/L adenine and 300 mg/L methionine). Spot test plates were grown at 30°C for 6 days before imaging with a Canon EOS 500D.

**Cloning and assembly for genome editing in *M. polymorpha***

CRISPR/Cas9 constructs were generated to target both the Mp*WDR1* and *2* loci with a single guide each and to target the Mp*WDR3* locus with two guides. Additionally, we used the *Csy4* CRISPR-associated ribonuclease system for polycistronic expression of four guides (two for Mp*WDR1* and two for Mp*WDR2*) or six guides (two for each of Mp*WDR1*, *2* and *3*). The constructs were assembled using parts from the OpenPlant kit (Sauret-Güeto *et al.*, 2020) along with some bespoke parts, as follows. Guides for direct cloning in L1_*lacZgRNA*-Ck2 or L1_*lacZgRNA*-Ck3 were annealed and integrated following Sauret-Güeto *et al.* (2020). The *Csy4* coding sequence was amplified by Phusion HF two-step PCR from pAC_06_L0_Csy4P2A with TK23 and 24, cloned in pUAP4 according to Sauret-Güeto *et al.* (2020) and assembled in an L1 construct with the *MpUBE2* promoter and the *Nos-35S* terminator, which was confirmed by restriction enzyme digestion and Sanger sequencing with TK71. Additionally, a bespoke version of the *MpUBE2* 5'UTR part was prepared with a 3' *Csy4* recognition sequence by Phusion HF two-step PCR with TK19-22 and cloned in pUAP4. Parts for the polycistronic guide constructs were amplified from this part and from pMODB2112 by Phusion HF two-step PCR with TK28-29,34-37,52-59 and assembled directly into pUAP4 using a bespoke SapI syntax and verified by Sanger sequencing. L2 and L3 assemblies were carried out according to Sauret-Güeto *et al*. (2020) and the final assemblies were confirmed by restriction enzyme digestion and Sanger sequencing (with p_CF, p_CR, AC188, TK41, TK45, TK49). The plasmids were introduced into *Agrobacterium tumefaciens GV3101* by electroporation and selected on LB medium with 25 µg/mL gentamicin, 50 µg/mL rifampicin and 50 µg/mL kanamycin or 100 µg/mL spectinomycin. Guide sequences are given in Table S1 and primer sequences in Table S2.

**Cloning and assembly for transcriptional reporters in *M. polymorpha***

The constructs were generated as described in Romani *et al.* (2024) except that for Mp*WDR1* and *2* a longer stretch, 3kb of upstream sequence, was used, in addition to the 5'UTR.

***M. polymorpha* spore transformation and genotyping**

Transformation of *M. polymorpha* sporelings, deriving from silica-dried archegoniophores carrying *Cam1x2* spores, was carried out according Annese *et al*. (2025), which was modified from Sauret-Güeto *et al.* (2020), itself based on Ishikazi *et al.* (2008). Sporelings were selected on half-strength Gamborg medium with 100 µg/mL cefotaxime and 0.5 µM chlorsulfuron and/or 20 µg/mL hygromycin. Primary transformants were transferred to new selective medium with the addition of 0.5% sucrose to encourage gemma cup production. Isogenic lines were established by culture of gemmae on selective medium, after which point lines were maintained on half-strength Gamborg. DNA extractions for genotyping were performed according to Frangedakis *et al*. (2020). Samples were stored at 4°C and 5 µL was used as the template for PCR amplification with KOD Xtreme Hot Start polymerase and primers TK80-81, 83-85, 90-94, 98, 99, 102, 103, 134-136. For gel assessment only, a positive plasmid template control and a negative water template control were included. For sequencing, the remaining

PCR product was purified with the GeneJet PCR purification kit, eluted in 20 µL dH2O and sent for Sanger sequencing. Primer sequences are given in Table S2.

##### *M. polymorpha* phenotyping

Induction of auronidin pigmentation was carried out according to Albert *et al.* (2018). 14-15 days after transfer to minimal medium, pigmentation status was scored visually and plants were imaged with a Keyence VHX Digital Microscope. Oil bodies were analysed in gemmae stained with 500 nM BODIPY 493/503 (Invitrogen). Gemmae were stained for 10 mins, rinsed twice in water with 0.1% Triton-X100 and imaged with a Leica SP5 Confocal Microscope, with excitation at 496 nm and 633 nm. Maximum intensity projections were prepared and merged for the Z-planes collected from the two channels using Fiji for ImageJ 2.9.0 (Schindelin *et al.*, 2012), and stained oil bodies were counted. In addition to oil bodies, staining was observed in rhizoid initials and slime papillae, and these structures were excluded from the count. The length of the perimeters of the gemmae were measured by Fiji on masks generated using thresholding (15-255) on maximum intensity projections of the Z-planes collected from the chlorophyll channel. Plotting and statistical analysis was carried out with the python packages *numpy*, *matplotlib*, *scipy*, *pandas* and *seaborn* using the Jupyter Notebook platform (Kluyver *et al.*, 2016; Virtanen *et al.*, 2020; Waskom, 2021).

#### Supplemental information

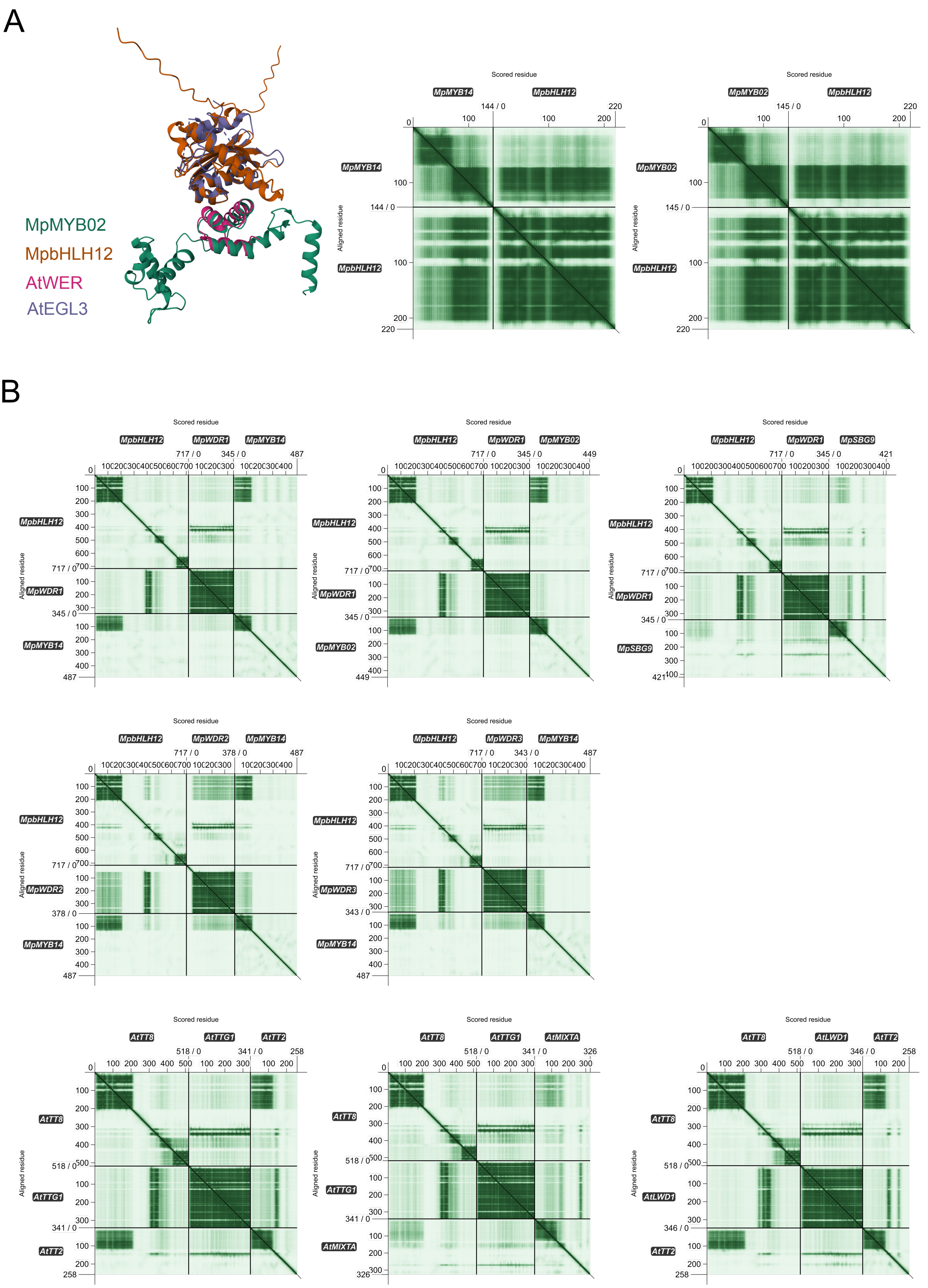


***Fig S1. Additional plots for the structural predictions***

***A.*** *Additional plots for the structural predictions for dimers from* M. polymorpha. *An AlphaFold Multimer dimer prediction for the MpbHLH12 N-terminal region and the R2R3MYB domain of MpMYB02 is aligned to the solved crystal structure of a dimer between the N-terminal region of AtEGL3 (aa 1-205) and the R3MYB repeat of AtWER (67-120)* (Wang *et al.*, 2022)*. The plots show the predicted aligned error (PAE) of the spatial position of each amino acid residue on the x-axis relative to each amino acid residue on the y-axis for each of the two* M. polymorpha *dimer predictions. A darker green colour in the plot indicates higher confidence (lower PAE) for the relative position of two residues (x,y). Each protein is aligned to itself and to each of the other proteins in the multimer prediction.* ***B.*** *Eight Alphafold Multimer trimer predictions were performed for combinations of proteins from* M. polymorpha *and* A. thaliana*. The predicted aligned error plots are given, as above. In each case, a bHLH, a WDR and a MYB protein were combined. Three combinations of proteins from* A. thaliana *are shown, representing a canonical MBW complex (in which AtTT8 is the bHLH member, AtTTG1 is the WDR member and AtTT2 is the MYB member), one where AtTT2 has been replaced by the paralogous MYB AtMIXTA, and one where AtTTG1 is replaced by the paralogous WDR AtLWD1. For* M. polymorpha*, five combinations are shown of the single IIIf bHLH with each of the two VIII-E MYBs or with the paralogous MYB MpSBG9 and each of the three TTG1-like WDRs.*

| BD | AD |
|---|---|
| MpbHLH12 | - |
| - | MpMYB02 |
| MpbHLH12 | MpMYB02 |
| MpWDR1 | - |
| - | MpbHLH12 |
| MpWDR1 | MpbHLH12 |

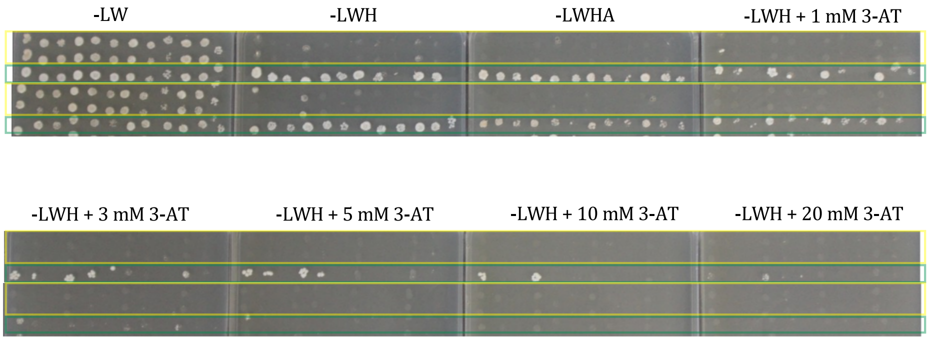


| BD | AD |
|---|---|
| MpbHLH12 | - |
| - | MpMYB14 |
| MpbHLH12 | MpMYB14 |
| MpWDR2 | - |
| - | MpbHLH12 |
| MpWDR2 | MpbHLH12 |

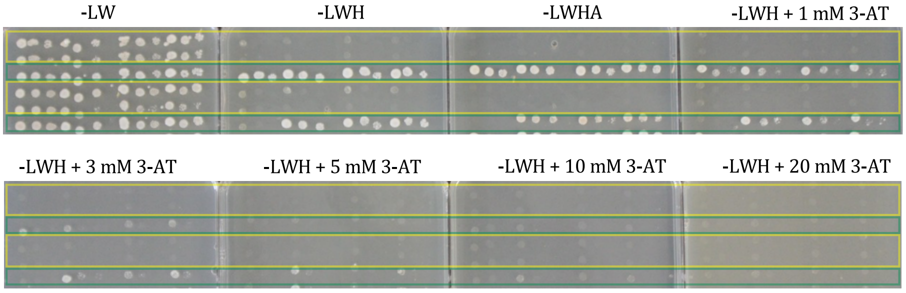


| BD | *PROM_Met17::* | AD |
|---|---|---|
| MpbHLH12 | MpWDR1 | mVenus |
| MpbHLH12 | MpWDR2 | mVenus |
| MpbHLH12 | - | mVenus |
| mVenus | - | MpMYB14 |
| MpbHLH12 | MpWDR1 | MpMYB14 |
| MpbHLH12 | MpWDR2 | MpMYB14 |
| MpbHLH12 | - | MpMYB14 |

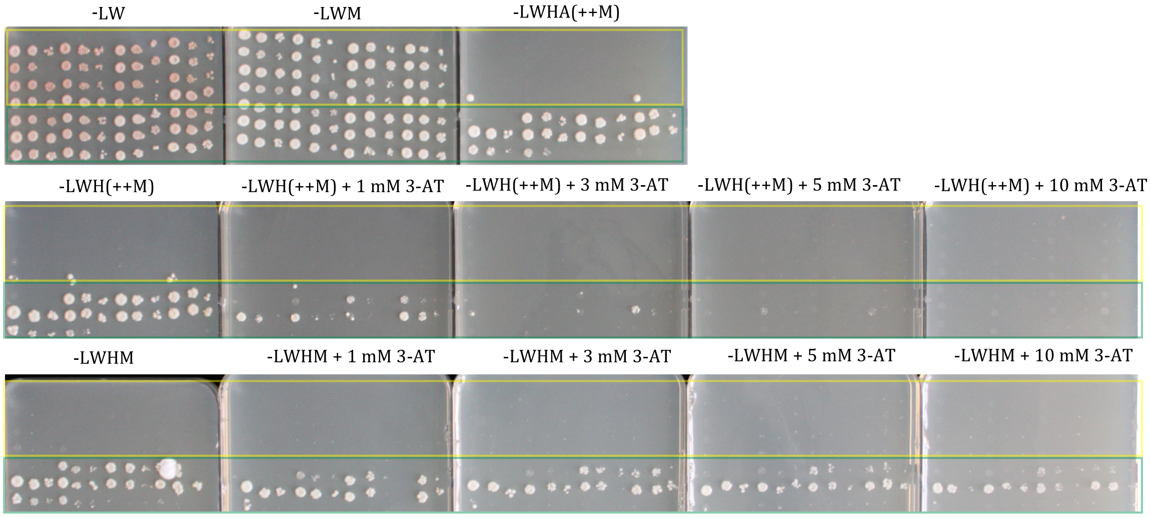


***Fig S2. Yeast spot test plates***

*Rows correspond to separate co-transformations. Autoactivation tests are highlighted in yellow and protein-protein interaction tests are highlighted in green. Four biological replicates were included, with two serial dilutions of each. Growth on medium without leucine and tryptophan (-LW) in each experiment confirms the colonies carry both plasmids with the genes of interests indicated in the tables in translational fusion with either the Binding Domain (BD) or Activation Domain (AD) of the yeast transcription factor GAL4. Protein-protein interaction between the proteins of interest allows a reconstituted GAL4 to drive expression of reporter genes from the GAL4 Upstream Activation Sequence (UAS). These reporter gene activities complement auxotrophic mutations in histidine and adenine biosynthesis genes in the background genotype, allowing the yeast to grow in the absence of these amino acids (-HA). 3-amino-1,2,4-triazole (3-AT) is a competitive inhibitor of the gene product of the histidine reporter. In one experiment, a* MET17 *cassette integrated on the BD plasmid drives methionine-repressible expression of a third gene.*

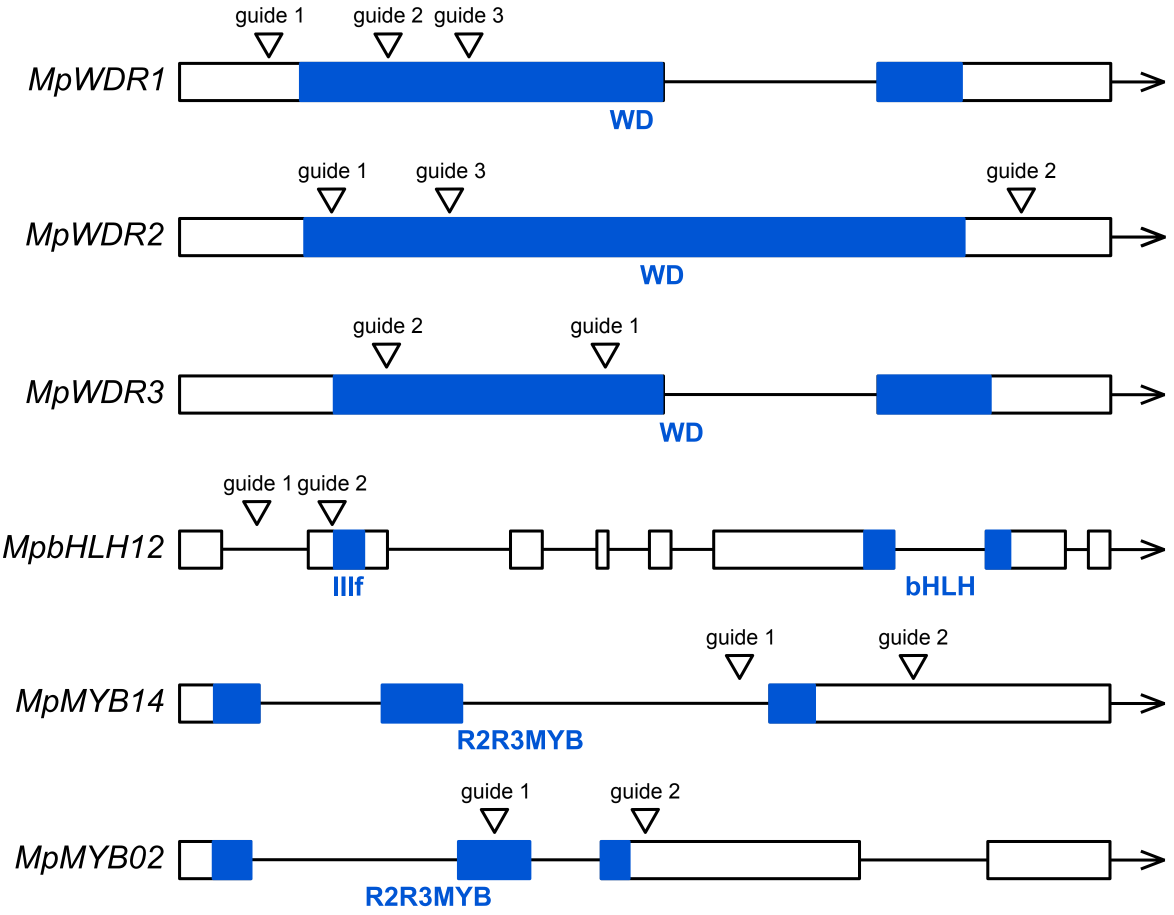


| **Line** | ***Edit(s)*** |
|---|---|
| *wdr1,2* line 20 | At the MpWDR1 locus, a 1 bp deletion at guide 3 and at the MpWDR2 locus, a 1 bp deletion at guide 3. |
| *wdr2* line 3 | *1 bp deletion at guide 3* |
| *wdr2* line 9 | *1 bp insertion at guide 3* |
| *wdr3* line 9 | *66 bp insertion at guide 1 and 2 bp deletion at guide 2* |
| *wdr3* line 11 | *1 bp deletion at guide 1 and 55 bp insertion at guide 2* |
| *bhlh12* line 1 | *1 bp deletion at guide 1* |
| *bhlh12* line 8 | *1 bp deletion and 1 bp insertion leading to frameshift in 400 bp region between guides 1 and 2* |
| *bhlh12* line 21 | *1 bp deletion at guide 1 and 52 bp deletion at guide 2* |
| *bHLH12_mut* line 34 | *In-frame deletion of entire region between guides 1 and 2* |
| *bhlh12* line 37 | *55 bp insertion at guide 2* |
| *myb14* line 16 | *4 bp insertion at guide 2* |
| *myb14* line 34 | *12 bp insertion at guide 1 and 1 bp deletion at guide 2* |
| *myb02* line 1 | *91 bp deletion at guide 1* |
| *myb02* line 3 | *1 bp insertion at guide 2* |

***Fig S3. CRISPR mutant lines overview***

*Gene models shown are not to scale. The positions of CRISPR/Cas9 guide target sequence are shown, and the mutations in the analysed lines are listed.*

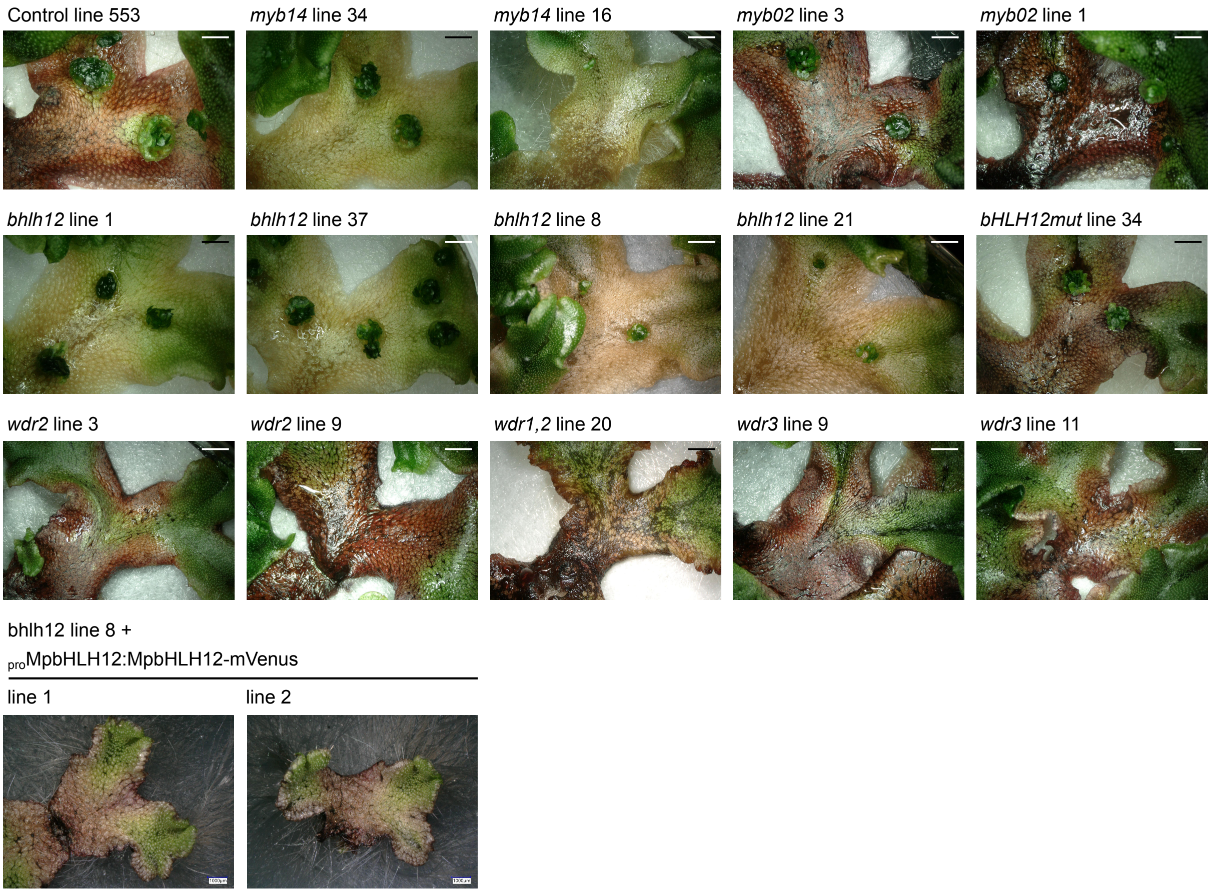


***Fig S4. Pigment scoring for additional independent lines***

*Two independent CRISPR/Cas9 loss of function mutant lines are shown for each of Mp*MYB14*, Mp*MYB02*, Mp*WDR2 *and Mp*WDR3*. A single line is shown for the control line and for the only* wdr1 wdr2 *double mutant line recovered. For Mp*bHLH12*, four loss of function lines are shown as well as one line with a mutation unlikely to impair protein function. Gemmae were germinated on ½ strength Gamborg and transferred to minimal medium at 4 weeks old. Plants were imaged 14-15 days later. Scale bars are 2 mm except for the complemented lines, for which they are 1 mm.*

***Table S1. CRISPR/Cas9 guide sequences***

| Target gene | Guide number | Guide sequence |
|---|---|---|
| *MpWDR1* | 1 | GCAGATCAATGAAGACAAGG |
| | 2 | GCTTTTGGCTCTCCTTGTCG |
| | 3 | CGAGGAGTACTCGAACAAGG |
| *MpWDR2* | 1 | GCTTCTGGCACTCCTTATCG |
| | 2 | GAGCTTGCTCCCGAAAGCAA |
| | 3 | AGAGGCTCTTGGCTTCCACG |
| *MpWDR3* | 1 | GTTGGAGTCCATGACAACAG |
| | 2 | GGACCTTCTGGCCACCACGG |
| *MpbHLH12* | 1 | TCCAGTTTCGAGATACACGG |
| | 2 | TTCCTCTTGACTGAGCTCGG |
| *MpMYB14* | 1 | CCTCGGAACACAGAGCACCG |
| | 2 | GCACTCCACCAGCTTCACGG |
| *MpMYB02* | 1 | CTCCGTCCAGACCTAAAGCG |
| | 2 | GCATCGTCCACTGAGCCAGG |

***Table S2. Other oligo sequences***

| | |
|---|---|
| MpbHLH12_Gateway_F (AC39) | CTACGACACGATGGCTGGAG |
| MpbHLH12_Gateway__R(AC40) | TTATCTGGAGCCCATCCCAGC |
| MpMYB14_Gateway__F (AC11) | ATGGTGCAACCGATACATTCG |
| MpMYB14_Gateway__R (AC12) | CTACCTCAGGCTGTGCATGTG |
| MpMYB02_Gateway__F (AC13) | ATGGGACGAGCTCCGTGTTG |
| MpMYB02_Gateway__R (AC14) | CTAGAAGGCAGTAGGGAGCTG |
| MpWDR1_Gateway__F (AC74) | ATGGCGAGCGAGAAGAAGGATG |
| MpWDR1_Gateway__R (AC75) | TCACACCCTGAGAATCTGCAAC |
| MpWDR2_Gateway__F (AC72) | ATGGACGGGTTCTCACAAGAACC |
| MpWDR2_Gateway__R (AC73) | TCACACTCGTAGAATTTGGAG |
| MpWDR3_Gateway__F (AC191) | ATGTCGAACCGAAATAGAACC |
| MpWDR3_Gateway__R (AC192) | TCATATTCTTAAAAGCTGCAGCTC |
| Gateway_attB1_adapter (AC87) | GGGGACAAGTTTGTACAAAAAAGCAGGCT |
| Gateway_attB2_adapter (AC88) | GGGGACCACTTTGTACAAGAAAGCTGGGT |
| Gateway_attL1 | TCGCGTTAACGCTAGCATGGATCTC |
| Gateway_attL2 | GTAACATCAGAGATTTTGAGACAC |
| MpWDR1_Gibson_F (TK9) | gtcagatacatagatacaattctattacccccatccatacATGGCGAGCGAGAAGAAGG |

| | |
|---|---|
| MpWDR1_Gibson_R (TK10) | gcgtgaatgtaagcgtgacataactaattacatgaTCACACCCTGAGAATCTGCAAC |
| MpWDR2_Gibson_F (TK11) | gtcagatacatagatacaattctattacccccatccatacATGGACGGGTTCTCACAAG |
| MpWDR2_Gibson_F (TK12) | gcgtgaatgtaagcgtgacataactaattacatgaTCACACTCGTAGAATTTGGAG |
| MpWDR3_Gibson_F (TK13) | gtcagatacatagatacaattctattacccccatccatacATGTCGAACCGAAATAGAAC |
| MpWDR3_Gibson_R (TK14) | gcgtgaatgtaagcgtgacataactaattacatgaTCATATTCTTAAAAGCTGCAGCTC |
| MET17_Gibson_F (TK15) | GCTGGTGGTGTTGTTGAATTGATCGCTTAAttcgaatcccttagctctca |
| MET17_Gibson_R (TK16) | gtatggatgggggtaatagaattg |
| CYC1_Gibson_F (TK17) | tcatgtaattagttatgtcacgctt |
| CYC1_Gibson_R (TK18) | gcaaattaaagccttcgagc |
| M13_F | TGTAAAACGACGGCCAGT |
| p_CF | gcaacgctctgtcatcgttac |
| p_CR | gtaacttaggacttgtgcgacatgtc |
| AC188 | GGAGCTTGCTCCCGAAAGC |
| TK19 | GTCTCaTACTgttagggaggtggggtggggttggg |
| TK20 | CTGCCTATACGGCAGTGAACgggcaccagcaccgccaagc |
| TK21 | gggGCTCTTCGTCTCaTACTgttagggaggtg |
| TK22 | gggGCTCTTCGTCTCaAAGCCTGCCTATACGGCAGTGAAC |
| TK23 | gggGCTCTTCGTCTCaAATGGACCACTACCTCGACATCAG |
| TK24 | gggGCTCTTCGTCTCaAAGCTCAGAACCACGGCACGAAGCC |
| TK28 | gggGCTCTTCGTCTCaAAGCCTGCCTATACGGCAGTGAACG |
| TK29 | gggGCTCTTCGTCTCaTACTgttagggaggtgg |
| TK34 | tttGCTCTTCgGCCAAAAGCCTGCCTATACGGCAGTGAACG |
| TK35 | gggGCTCTTCgGGCTCTCCTTGTCGGTTTTAGAGCTAGAAATAGCAAGT |
| TK36 | tttGCTCTTCgGGGAGCAAGCTCCTGCCTATACGGCAGTGAACG |
| TK37 | gggGCTCTTCgCCCGAAAGCAAGTTTTAGAGCTAGAAATAGCAAGTTAAA |
| TK41 | TAAAACCCGTGGTGGCCAGAAGGTCC |
| TK45 | TAAAACCCGAGCTCAGTCAAGAGGAA |
| TK49 | TAAAACCCGTGAAGCTGGTGGAGTGC |
| TK50 | GCTAGATTCGTCTCCAAG |
| TK52 | gggGCTCTTCgTTGATCTGCCTGCCTATACGGCAGTGAAC |
| TK53 | gggGCTCTTCgCAATGAAGACAAGGGTTTTAGAGCTAGAAATAGCAAGT |
| TK54 | gggGCTCTTCgATAAGGAGTGCCAGAAGCCTGCCTATACGGCAGTGAACG |
| TK55 | gggGCTCTTCgTATCGgttttagagctagaaatagcaagttaaaataaggct |
| TK56 | gggGCTCTTCgAACCTGCCTATACGGCAGTGAACG |
| TK57 | gggGCTCTTCgGTTGGAGTCCATGACAACAGGTTTTAGAGCTAGAAATAGCAAG |
| TK58 | tttGCTCTTCtCGTGGTGGCCAGAAGGTCCCTGCCTATACGGCAGTGAACG |
| TK59 | gggGCTCTTCgACGGGTTTTAGAGCTAGAAATAGCAAGT |

| | |
|---|---|
| TK71 | ATCGGGGAACCAAAGACAACTG |
| MpWDR1_genotyping_F (TK80) | ATGGCGAGCGAGAAGAAGGATG |
| MpWDR1_genotyping_R (TK81) | TCTCACCTGATGGCGTTGTAGC |
| MpWDR2_genotyping_F (TK136)__ | AAGGTGGAGATTGTGCAGCTGG |
| MpWDR2_genotyping_R (TK83) | CTTCACACACTCTGGCTCCACTG |
| MpWDR3_genotyping_F (TK84) | ACGCGCTGAATTGGACTATCAGG |
| MpWDR3_genotyping_R (TK85) | TCACCTGATGTCGCTGCAGATC |
| Csy4_genotyping_F (TK90) | ACCACTACCTCGACATCAGGCTC |
| Csy4_genotyping_R (TK91) | TGAGCCTCTCTGGGTTGGACTTG |
| Cas9_genotyping_F (TK92) | ctacgtgggacctctcgctagag |
| Cas9_genotyping_R (TK93) | agccttccccaaccagtgtatcttc |
| MpbHLH12_genotyping_F (TK98) | GATTTGTGGGCTCTTCTTGG |
| MpbHLH12_genotyping_R (TK99) | ATGCATAGCAGATAGAACCACTC |
| MpMYB14_genotyping_F (TK102) | CACGTTTCTCGTTCTGTGCAC |
| MpMYB14_genotyping_R (TK103) | CTCACTTGGTCGAAGCTGTACTC |
| MpMYB02_genotyping_F (TK134) | GTGAACCGTCTCAGTGTGAAGC |
| MpMYB02_genotyping_R (TK135) | CATTCAATTCGGCCTGATGAG |

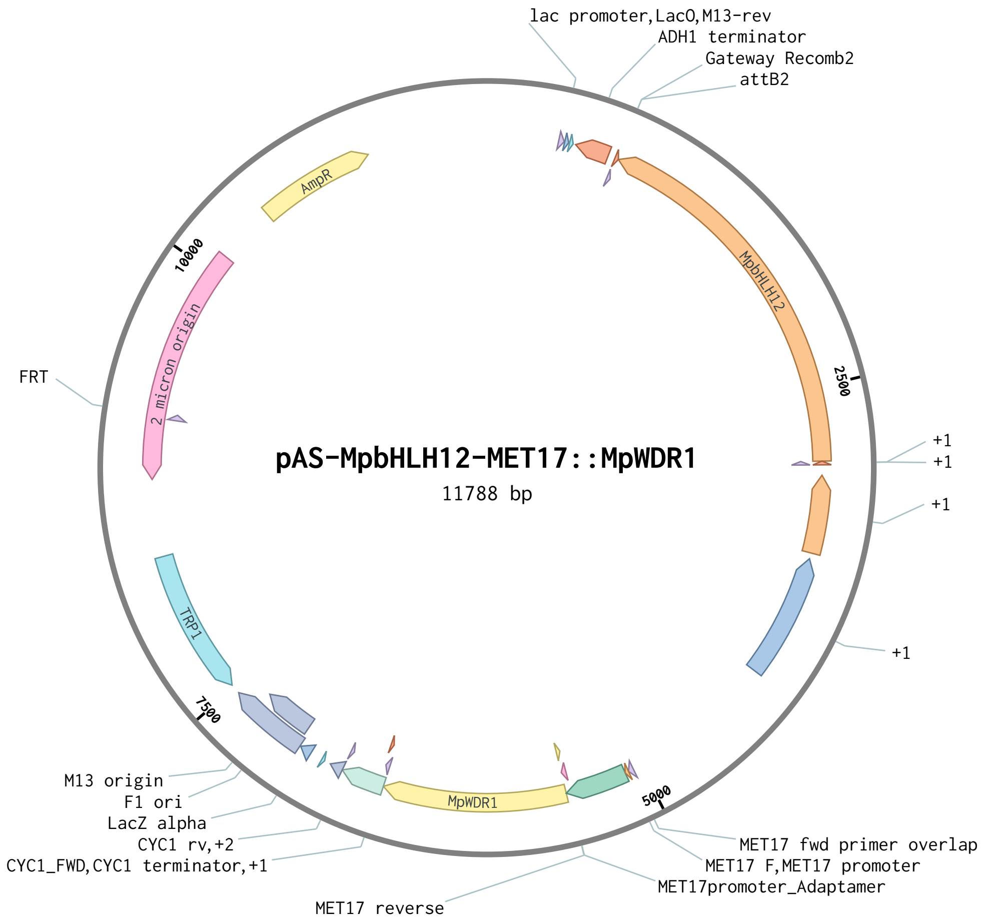


***Fig S5. Bespoke plasmid for tripartite interaction tests in*** **Saccharomyces cerevisiae.**

*The construct with Mp*WDR1 *is shown, the same constructs were made with Mp*WDR2 *or Mp*WDR3 *in its place.*

### Additional files

‘raw_counts.csv’: Raw oil body counts, perimeter lengths and area measurements
